# Climate drives spatial variation in Zika epidemics in Latin America

**DOI:** 10.1101/454454

**Authors:** Mallory Harris, Jamie M. Caldwell, Erin A. Mordecai

## Abstract

Between 2015 and 2017, Zika virus spread rapidly through populations in the Americas with no prior exposure to the disease. Although climate is a known determinant of many *Aedes*-transmitted diseases, it is currently unclear whether climate was a major driver the of Zika epidemic and how climate might have differentially impacted outbreak intensity across locations within Latin America. Here, we estimated force of infection for Zika over time and across provinces in Latin America using a time-varying Susceptible Infectious Recovered model. Climate factors explained less than 5% of the variation in weekly transmission intensity in a spatiotemporal model of force of infection by province over time, suggesting that week to week transmission within provinces may be too stochastic to predict. By contrast, climate and population factors were highly predictive of spatial variation in the presence and intensity of Zika transmission among provinces, with pseudo R^2^ values between 0.33 and 0.60. Temperature, temperature range, rainfall, and population size were the most important predictors of where Zika transmission occurred, while rainfall, relative humidity, and a nonlinear effect of temperature were the best predictors of Zika intensity and burden. Surprisingly, force of infection was greatest in locations with temperatures near 24°C, much lower than previous estimates from mechanistic models, potentially suggesting that existing vector control programs and/or prior exposure to other mosquito-borne diseases may have limited transmission in locations most suitable for *Aedes aegypti*, the main vector of Zika, dengue, and chikungunya viruses in Latin America.

## Introduction

The emergence and re-emergence of mosquito-borne diseases presents a global public health concern, yet trends in mosquito-borne disease transmission are hard to predict because they are influenced by the underlying immunity in the population, which is usually unknown [1– 3]. The emergence and spread of Zika virus through Latin America in a population with no prior exposure or immunity to the disease provides an opportunity to characterize relationships among the environment, populations, and disease transmission without the confounding effects of pre-existing immunity (although cross-reactivity between dengue antibodies and Zika virus may provide some protection) [4]. Between 2015 and 2017, Zika rapidly spread to 51 countries and territories in the Americas, with over 800,000 cases reported [5]. Here, we quantified the force of infection for Zika and studied factors that contributed to variation in transmission across time and space as Zika emerged.

Force of infection, or the per capita rate at which susceptible individuals contract an infection, can be used to compare disease transmission over the course of an outbreak and across geographic regions [2,6]. Force of infection estimates can account for variation in the number of immune individuals and entomological factors that influence the time it takes for mosquito-borne diseases to be transmitted [2,6]. It can take up to several weeks for mosquitoes to spread diseases because transmission requires a mosquito to bite an infectious person, the pathogen to replicate and disseminate in the mosquito (during the extrinsic incubation period), and the mosquito to bite and infect a susceptible person [2,7–9]. Force of infection can be calculated over time to study how transmission rate changes over the course of an outbreak and to compare outbreaks of the same disease in different places [2,6,10–12].

The progression of an epidemic can be characterized and compared through measures of burden, intensity, and transmission rate [13,14]. We can quantitatively differentiate between epidemics with sharp peaks in case incidence that quickly burn through the susceptible population and more sustained, low-level transmission [6,15,16]. Instances when high case incidence coincides with a low force of infection may indicate the presence of imported cases [17]. Each of these epidemic metrics may be affected to a different extent by climate and population factors, in addition to country and province-level features like existing transportation infrastructure and vector control resources [4,9,18,19]. Additionally, conditions that enable the establishment of a disease may differ from those that drive higher epidemic intensity following establishment [20].

Climate is a key determinant for whether mosquito-borne disease transmission can occur and the intensity of outbreaks because of its effects on vector dynamics. The primary vector of Zika in Latin America is *Aedes aegypti*, which also spreads chikungunya and dengue viruses [21]. Disease transmission can only occur if climate conditions are suitable for pathogen and vector survival and reproduction, while the intensity of transmission may be affected by how close humidity, temperature, and rainfall are to optimal climate conditions for vector and pathogen performance [2,9,22–27]. Humidity is positively associated with mosquito survival [23]. Temperature influences mosquito fecundity, development, and survival, as well as factors contributing directly to transmission, including the extrinsic incubation period and mosquito biting rate [9,24,27–29]. These thermal responses are nonlinear with optimal disease transmission expected at 29°C for viruses transmitted by *A. aegypti* based on mechanistic models parameterized from laboratory experiments [20,25]. Mosquitoes require water to complete their life cycle and breed, but the relationship between mosquito abundance and rainfall is variable [30–36]. The most notable reasons for this varying relationship are associated with extreme conditions, such as heavy rainfall that can kill and wash away larvae [37,38], and drought, which in certain regions promotes humans storing water in open containers that serve as breeding habitats [35,36]. Understanding how climate drives the emergence and intensity of Zika will be important for identifying regions where the disease is likely to become endemic, and, more generally, for predicting the potential trajectories of other mosquito-borne diseases that could emerge and re-emerge in the future, such as o’nyong nyong, Mayaro, and yellow fever [1,19,39– 41].

In this study, we estimate weekly force of infection for Zika from human case reports across Latin America to examine the role of climate in driving the emergence and intensity of the 2015 to 2017 outbreak. Specifically, we use the models to ask how climate and population variation affect (1) when and where epidemics occur, (2) epidemic dynamics over time, and (3) geographic variation in the intensity of epidemics. We use disease case reports and force of infection estimates in two modeling frameworks. First, we examine variation in force of infection over time within provinces to understand how strongly climate predicts the probability of weekly local transmission and the intensity of weekly force of infection. Then, we examine spatial variation in several epidemic metrics, including total human cases and mean force of infection, to understand how climate and population factors shape epidemics geographically.

## Materials and Methods

### Epidemiological Data

To investigate Zika transmission dynamics over time and space in Latin America, we downloaded and preprocessed publicly available human case data. We used weekly suspected and confirmed Zika cases between November 2015 and November 2017 for 156 provinces across seven countries in Latin America (Colombia = 32 provinces, Dominican Republic = 32 provinces, Ecuador = 24 provinces, El Salvador = 14 provinces, Guatemala = 22 provinces, and Mexico = 32 provinces) from the Centers for Disease Control and Prevention (CDC) Zika Data Repository, which includes epidemiological bulletins provided by each country’s ministry of health [42]. We excluded fourteen provinces with fewer than ten weeks of case reporting or irregular reporting intervals, because those provinces provided insufficient data to observe meaningful trends in transmission (excluded provinces: Ecuador = eight provinces, Guatemala = two provinces, and Mexico = four provinces). For the remaining provinces, we temporally interpolated case data for weeks with missing data and reporting errors by averaging cases from the weeks immediately preceding and following these intervals.

### Weather Data

To investigate the effects of climate on Zika transmission, we downloaded weather data and calculated climate metrics with time lags relevant to diseases spread by the *Aedes aegypti* vector. We downloaded daily mean relative humidity, total rainfall, and mean, minimum, and maximum temperatures from Weather Underground [43]. For each province, we used the weather station nearest to the province’s centroid that had the most complete climate record in the timespan corresponding to the case data. We excluded from our analyses an additional fifteen provinces that had no nearby weather station reporting in the desired time period (excluded provinces: Colombia = six provinces, Dominican Republic = one province, Ecuador = two provinces, El Salvador = two provinces, Guatemala = two provinces, and Mexico = two provinces), and further excluded 277 weeks with mean temperatures outside of the range of 0-40°C and rainfall values exceeding 250mm, as these extreme values are likely to be weather station errors. Our analyses included the remaining 127 provinces with a total of 7,109 weekly observations of epidemiological and weather data. We believe weather station data provides more accurate measurements of climate near populated areas (as weather stations are located at airports or operated for personal use) compared with modeled weather data such as the NOAA National Centers for Environmental Prediction Reanalysis data (NCEP; see Figure S1 for a comparison between data from Weather Underground and NCEP, a gridded global model based on satellite data), and therefore chose to conduct our analyses with weather station data with a reduced sample size due to missing weather stations in some provinces.

To investigate the spatiotemporal dynamics of transmission, we calculated lagged climate metrics (see Figure S2 for heatmaps showing variation in climate by province over time), as humidity, temperature, and rainfall influence transmission at a delay, which is commonly assumed to be between one and two months [30–32,34,35,44]. Specifically, we calculated humidity, mean temperature, and temperature range (difference between the maximum and minimum temperatures observed) over a three-week period, lagged by six weeks from the week of case reporting (i.e., nine to seven weeks prior, following previous work) [20,45,46]. Similarly, we calculated the cumulative rainfall over a six-week period, lagged by three weeks from the week of case reporting (i.e., nine to four weeks prior), extending the window applied to lagged temperature period by three weeks in order to better capture the effects of water accumulation over time [47,48]. To compare differences in overall epidemic characteristics (e.g., total number of cases, mean force of infection Table 1; Figure 1) among provinces, we also calculated province-level mean humidity, mean temperature, temperature range, and cumulative rainfall over the biologically relevant time lag described above.

**Table 1:** Response variables for the spatiotemporal (ST) and spatial (S) models, with their corresponding thermal optima. R^2^ values were calculated for the spatiotemporal models and pseudo R^2^ values were calculated for the spatial models using Nakagawa’s pseudo R^2^ for the mixed effect local transmission model (which produces a marginal and conditional pseudo R^2^) and Nagelkerke’s pseudo R^2^ for the other spatial models. For the best fit models, temperature and humidity correspond to mean temperature and mean humidity over a three-week period, lagged by six weeks from the week of case reporting. Temperature^2^ corresponds to squared temperature over a three-week period, lagged by six weeks from the week of case reporting. Rainfall corresponds to cumulative rainfall over a six-week period, lagged by three weeks from the week of case reporting. In the case of the spatial models, climate covariates are averaged over the course of the epidemiological data and city pop corresponds to the population of each province’s largest city.

| Response variable | Description | Model Type | Thermal Optimum (°C) | R<sup>2</sup>/ Pseudo R<sup>2</sup> | Sample Size | Best Fit Model |
| --- | --- | --- | --- | --- | --- | --- |
| Weekly Local Transmission | The likelihood that $\beta_t$ exceeded zero for a given week | ST | N/A | 0.037 | 7,109 | $P(\beta_t > 0) \sim$<br>$\sim 0.07 \times \text{temperature}$<br>$+ 0.06 \times \text{humidity}$ |
| Intensity of Weekly Force of Infection | The magnitude of $\ln(\beta_t)$, given that it exceeded one | ST | N/A | 0.011 | 2,031 | $\ln(\beta_t) \sim$<br>$0.07 \times \text{temperature}$<br>$- 0.08 \times \text{humidity}$<br>$+ 0.04 \times \text{rainfall}$ |
| Local Transmission | The likelihood that $\beta_t$ exceeded zero for any week | S | N/A | Marginal: 0.281<br>Conditional: 0.599 | 127 | $P(\text{local trans}) \sim$<br>$0.49 \times \text{temperature}$<br>$+ 1.15 \times \text{humidity}$<br>$- 0.07 \times \text{rainfall}$<br>$+ 0.82 \times \text{city pop}$ |
| Maximum Force of Infection | Logged maximum value for $\beta_t$ in provinces with local transmission | S | 23.93<br>(95% CI: 22.02-26.90) | 0.534 | 114 | $\ln(\max(\beta_t)) \sim$<br>$- 2.30 \times \text{temperature}^2$<br>$+ 2.44 \times \text{temperature}$ |
| Mean Force of Infection | Logged mean value for $\beta_t$ in provinces with local transmission | S | 23.63<br>(95% CI: 22.20-25.49) | 0.596 | 114 | $\ln(\text{mean}(\beta_t)) \sim$<br>$- 2.78 \times \text{temperature}^2$<br>$+ 2.75 \times \text{temperature}$<br>$+ 0.19 \times \text{city pop}$ |
| Cumulative Cases | Total number of cases reported | S | 22.51<br>(95% CI: 20.57-30.68) | 0.330 | 127 | $\text{cumulative cases} \sim$<br>$- 1.68 \times \text{temperature}^2$<br>$+ 1.69 \times \text{temperature}$<br>$+ 0.16 \times \text{humidity}$<br>$+ 0.21 \times \text{city pop}$ |
| Mean Weekly Cases | Mean number of cases in weeks with any cases reported | S | 23.69<br>(95% conf: 19.57-26.12) | 0.348 | 127 | $\text{mean weekly cases} \sim$<br>$- 1.59 \times \text{temperature}^2$<br>$+ 1.69 \times \text{temperature}$<br>$+ 0.14 \times \text{rainfall}$<br>$+ 0.35 \times \text{city pop}$ |

**Figure 1:**
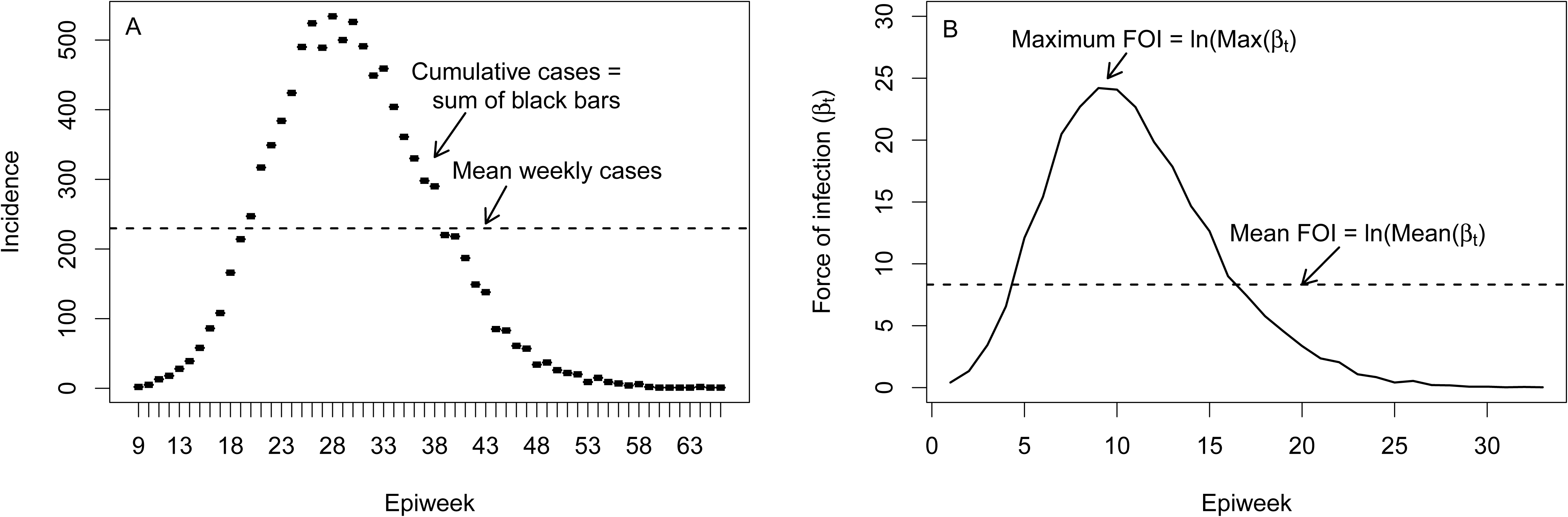
Illustration of epidemic metrics that characterize the presence, burden, and intensity of outbreaks. (A) Cartoon of weekly incidence during an outbreak where cumulative cases are calculated by taking the sum of all cases across all weeks during the outbreak, and mean cases (dashed horizontal line) are calculated as the mean of all cases in weeks with at least one case during the outbreak. (B) Cartoon of logged weekly force of infection during an outbreak where the highest value corresponds to maximum force of infection and the dashed horizontal line corresponds to the mean force of infection.

### tSIR Model

We estimated weekly force of infection by fitting time-varying Susceptible Infectious Recovered (tSIR) models to the human case data [42]. We assumed a static population size because the human population did not change significantly over the course of the epidemic, and we assumed that Zika was introduced to a fully susceptible population since it had never been documented before in this region. Therefore, we set the initial susceptible population (*S*_0_) as the most recent estimate of population (*N*) available from *City Population* as of June 2017, and modeled the susceptible population following Zika introduction as *S*_*t*_ = *S*_*t*−1_ − *I*_*t*_ where *S* indicates the susceptible population at time (*t*) or (*t* − 1) and *I* indicates the infected population at time (*t*) [49]. We defined effective infectious individuals based on the modification of the tSIR model for vector-borne diseases developed by Perkins et al. 2015 for chikungunya transmission because of similarities in the transmission ecology of their shared vector [2,34]. Using the infectious periods derived for chikungunya, we assumed that humans can transmit the parasite to mosquitoes for five days after reporting symptoms and that secondary human infections can arise from primary cases reported one to five weeks prior [50]. We calculated the effective number of infectious individuals 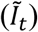 as a weighted sum of infectious individuals (*I*_*t*−*n*_) within a five-week serial interval (k=5). We used the same methods as the Perkins et al. 2015 model for chikungunya transmission to solve for the weights (*ω*_*n*_) and substituted the appropriate serial interval distribution for Zika (a Gaussian distribution with mean 20 days and standard deviation 7.4 days) [2,51,52]:

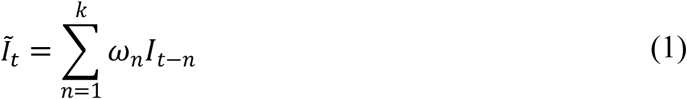

Individuals entered the recovered class after the five-week serial interval. We fit tSIR models to case data for each province to calculate time-varying force of infection (*β*_*t*_):

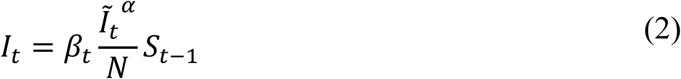

Here, the mixing parameter *α* indicates homogeneous frequency-dependent contact rate between cases in each province. We fixed the mixing parameter due to the tradeoff between capturing heterogeneity in transmission or mixing, a method commonly used in tSIR studies focused on transmission dynamics [10,53–56]. Given the similar transmission dynamics of chikungunya and Zika, we set the mixing parameter equal to 0.74, the value for chikungunya calculated in the Perkins et al. study [2,51]. By rearranging equation 2, we solved for ln(*β*_*t*_) using the following equation:

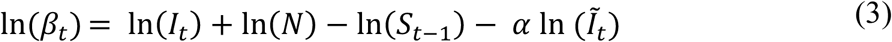

### Spatiotemporal Model

We investigated how strongly climate predicts the probability of weekly local transmission and the intensity of weekly force of infection by regressing force of infection against biologically relevant lagged and normalized climate factors. Since the transmission parameters (*β*_*t*_) were zero-inflated, we fit a two-step hurdle model that first predicts the probability that the force of infection, *β*_*it*_, will exceed zero in a given province (*i*) and week (*t*) (i.e., presence of local transmission), and then predicts the magnitude of logged force of infection, ln (*β*_*it*_) (i.e., intensity), given that *β*_*t*_ exceeds zero. To capture nonlinear relationships between transmission and temperature, we included a squared term for temperature [2,9,20].

We performed stepwise model selection using backward elimination, selecting the best fit models for both steps of the hurdle model as those with the highest adjusted R^2^, and further constraining the selection process to prohibit models that include a nonlinear temperature term without a linear temperature term. To account for autocorrelation between observations within each province over time, we fit panel models with a two-way random effect:

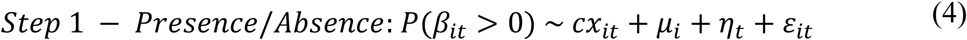

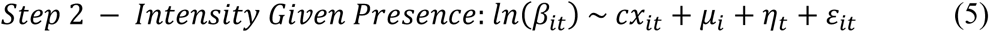

The hurdle models include climate covariates (*x*), their corresponding coefficients (*c*), time-independent province random effect (µ), province-independent random effect of week (η), and an observation-specific error term (ε) accounting for time (*t*) and province (*i*). Regressions were conducted in R statistical software version 3.4.3 using the plm function in the plm package [57]. We also fit alternative fixed effect spatiotemporal models with month or month nested in province fixed effects (Table S3-4).

### Spatial Models

We examined the factors that drove broad-scale geographic differences in Zika transmission by using linear regressions to compare epidemic metrics with normalized climate and population factors. The response variables included epidemic metrics based on incidence and force of infection (presence of local transmission, logged maximum force of infection, logged average force of infection, cumulative cases, and mean weekly cases), where each response variable was aggregated to a single value per province (Table 1; Figure 1). To account for additional variation among countries beyond climate and population factors, we included country as a fixed effect. Since some countries had local transmission in all provinces, we included country as a random effect in the spatial model for the probability of local transmission in a given province. For the population factor, we used the number of people living in the largest city in each province based on census results and official estimates as of June 2017 downloaded from *City Population* [49]. The mean value of each lagged climate covariate was taken for each province across the time that cases were reported. For each spatial model, we performed stepwise model selection using backwards elimination, selecting the best fit models as those with the lowest AIC, and calculated pseudo R^2^ with Nakagawa and Shielzeth’s method for the model of the probability of local transmission and Cragg, Uhler, and Nagelkerke’s method for remaining spatial models [58–61]. All regressions were conducted in R statistical software version 3.4.3 using the glm function in the base package.

## Results

### Epidemiological Data and tSIR Model

The duration and burden of Zika transmission and timing of epidemic peaks varied considerably both between and within countries (Figure 2). Within the countries included in this study, Zika emerged earliest in El Salvador in November of 2015, and latest in Colombia, which first reported a Zika case almost one year later in January of 2016 (Figure 2A). The weights for the serial interval were: *ω*_1_ = 0.081, *ω*_2_ = 0.448; *ω*_3_ = 0.372; *ω*_4_ = 0.089; *ω*_5_ = 0.010. Force of infection ranged from 0 to 15.409 cases per 100,000 people per week (model fit shown in Figure S3). In all countries there were weeks with no weekly local transmission (*β*_*t*_ = 0), but high case incidence resulting from imported cases. All countries included in our analyses had at least one province with local transmission (Figure 2B).

**Figure 2:**
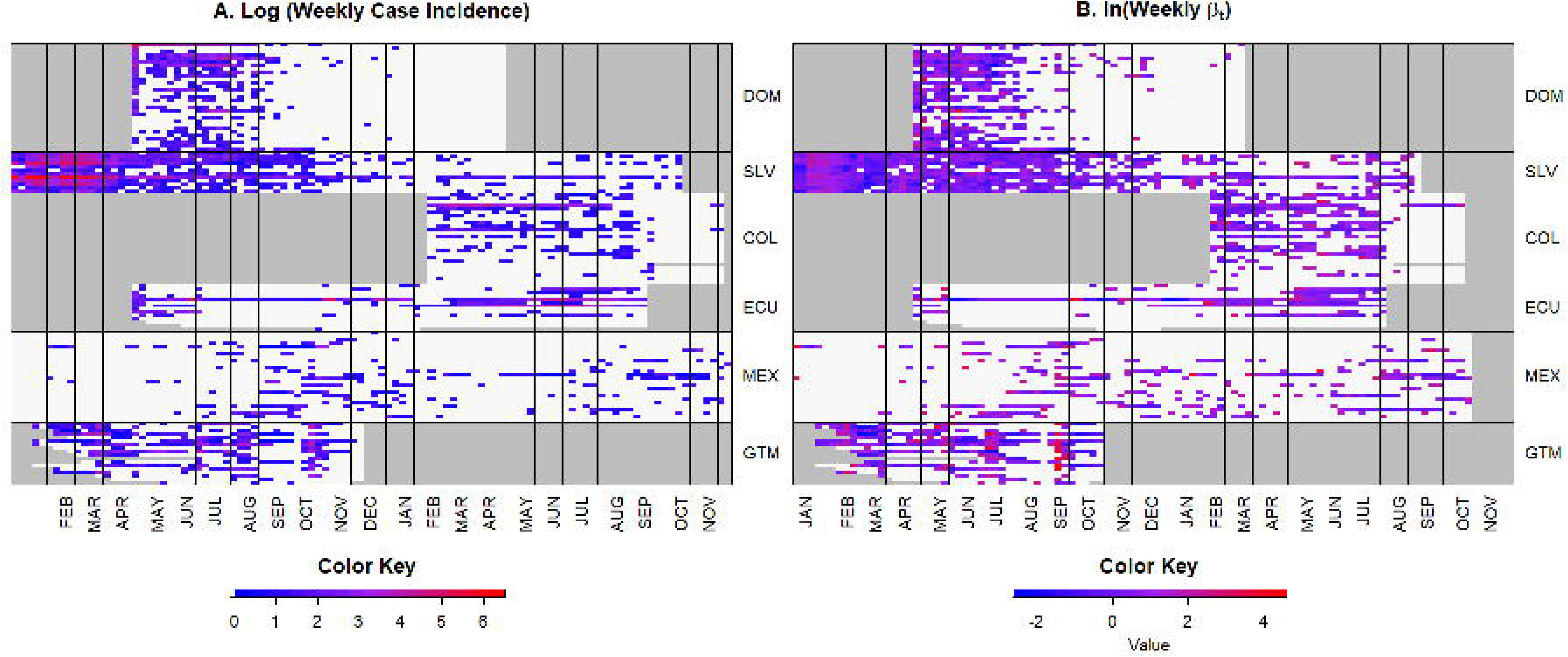
Heatmaps of weekly case incidence and force of infection. Each row corresponds to a single province over time, with lines separating the following countries: DOM = Dominican Republic, SLV = El Salvador, COL = Colombia, ECU = Ecuador, MEX = Mexico, GTM = Guatemala. Gray shading indicates no available data. The color represents (A) weekly logged cases for each province and (B) logged force of infection. Red indicates high values and blue indicates low values. White indicates (A) no cases reported or (B) no presence of local transmission (*β*_*t*_=0) in a given week.

### Spatiotemporal Model

The spatiotemporal hurdle model indicated that different climate factors were associated with the presence of weekly local transmission versus the intensity of weekly force of infection in a given province (Tables 1, S1-2). The best fit model for the presence of weekly local transmission included positive linear relationships with mean temperature and humidity and explained 3.7% of the variation in the likelihood of week to week transmission within provinces. The best fit model for the intensity of weekly force of infection included negative linear effects of mean temperature and humidity, as well as a positive linear effect of rainfall. This model explained 1.1% of the variation in the intensity of weekly force of infection. Alternative fixed effects models with month or month nested in province fixed effects produced similar results (Table S3-4).

### Spatial Models

The spatial models indicated that geographic variation in epidemics was driven by climate and human population sizes (Figure 4, Tables S5-9). There was considerable geographic variation in Zika presence and intensity within and among countries in Latin America (e.g., Figure 3). We found a positive linear relationship between the presence of local transmission and humidity and mean temperature (Figure 4). For all other epidemic metrics, we found a unimodal relationship with mean temperature, although this relationship is not statistically significant in the models of cumulative cases and mean weekly cases (positive linear effect of mean temperature and negative quadratic effect of mean temperature; Figures 4-5). In provinces that had local transmission, mean force of infection peaked at 23.63°C (95% confidence interval: 22.20°C-25.49°C; Figure 5B). The population of the largest city had a positive relationship with all epidemic characteristics aside from maximum force of infection (Figure 4). Mean humidity had a slight positive linear relationship with cumulative cases, while rainfall had a slight positive relationship with average weekly cases (Figure 4). All spatial models fit well based on their pseudo R^2^ values (0.33-0.60) (Table 1).

**Figure 3:**
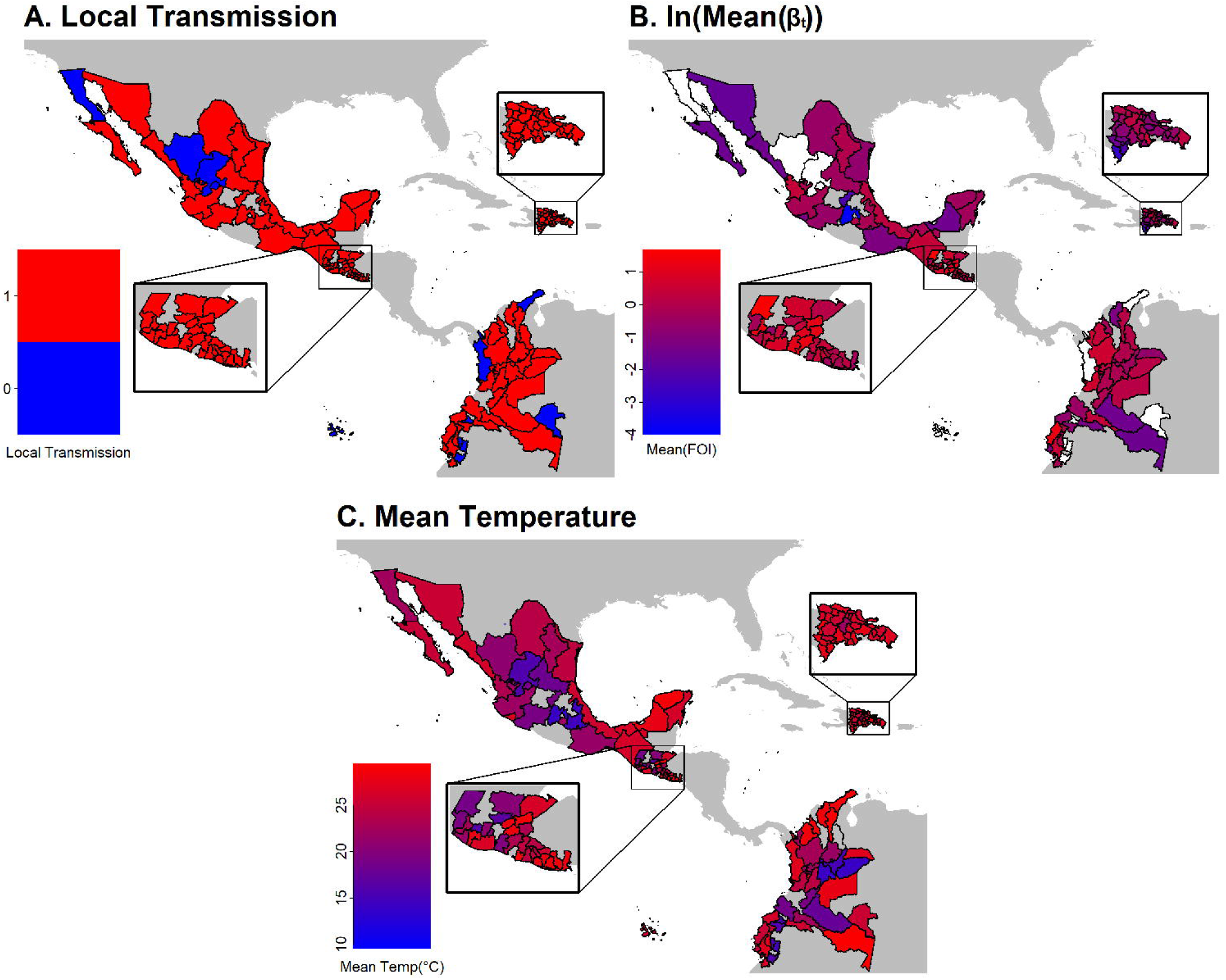
Geographic differences in epidemic metrics. Provinces are colored by (A) whether local Zika transmission was present (red) or absent (blue); (B) logged mean force of infection; and (C) mean temperature. Provinces that we excluded from this study due to lack of epidemiological or climate data (see Methods) are white. Countries not included in this study are grey.

**Figure 4:**
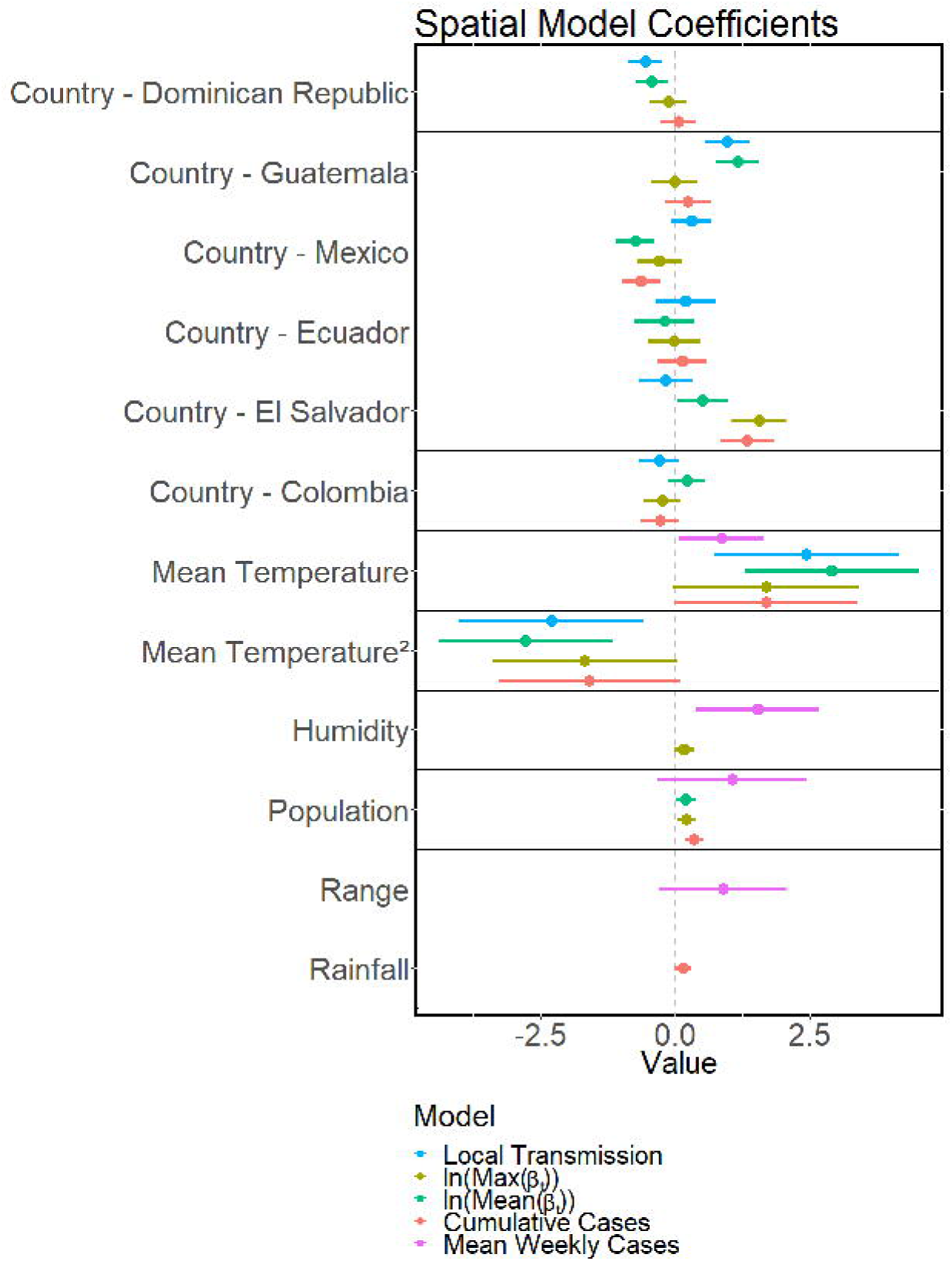
Coefficients for factors driving geographic differences in epidemics. Plots display coefficients included in the best fit spatial model for presence local transmission (lavender), maximum force of infection (blue), mean force of infection (green), cumulative cases (olive), and mean weekly cases (salmon). Points and horizontal lines indicate the mean coefficient values and 95% confidence intervals, respectively. Note that the presence of local transmission model is excluded from the panels reporting country fixed effects, as this model incorporates a random effect of country.

**Figure 5:**
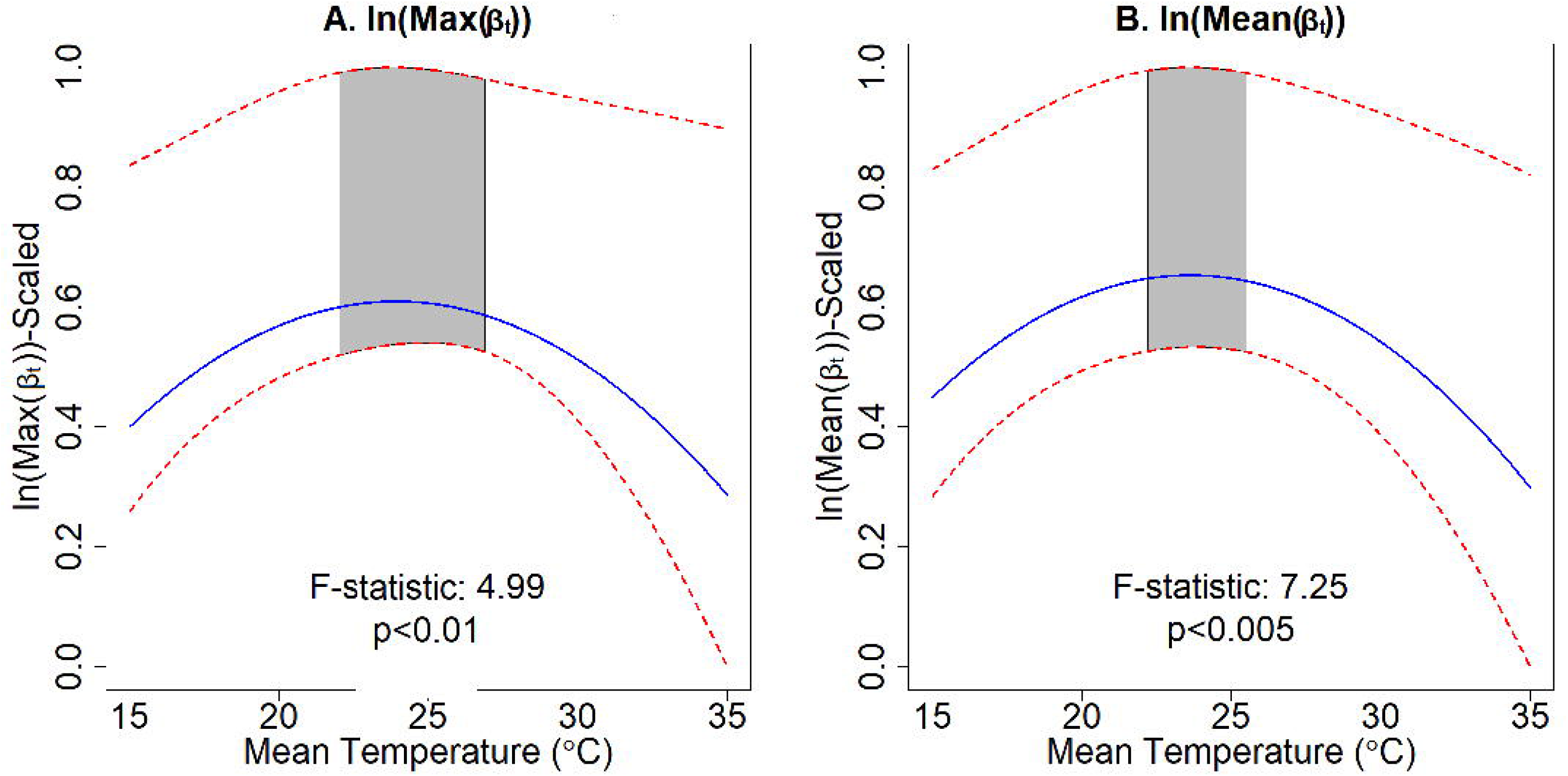
Nonlinear relationship between temperature and epidemic characteristics. Plots showing the relationship between mean temperature and (A) logged maximum force of infection and (B) logged mean force of infection. The model prediction is indicated by the blue lines, and the 95% confidence interval is indicated by the red dashed lines. The gray shaded region indicates the 95% confidence interval for the thermal optimum. The F-statistic for the nonlinear effect of temperature and its p-value are given in the inset. The response variables are scaled to be between zero and one.

## Discussion

Climate and population factors were strong drivers of geographic variation in Zika epidemics, with different factors influencing where local transmission occurred compared with the intensity of transmission. Relative humidity, temperature, temperature range, and the population of the largest city were key determinants for whether local transmission occurred, and these factors could be used to determine regional likelihood for future *Aedes aegypti* transmitted diseases. Both the spatiotemporal and spatial models for the presence of local transmission included a positive linear effect of temperature, while the burden and intensity of epidemics (based on the other spatial models) included nonlinear effects of temperature (Table 1, Figure 4A). The large variation in epidemic characteristics across countries (Figures 2, 3A-B, 4) suggests that additional factors beyond climate and population size may also be important in driving Zika transmission. Understanding how climate drives different epidemic metrics is valuable for preparing for and responding to outbreaks. For example, the infrastructure and resources needed to respond to a large outbreak that spreads rapidly through the population, like what occurred in the Dominican Republic and El Salvador, are very different from what is needed to respond to an outbreak that affects fewer people at one time but lasts many months, as in Colombia and Guatemala.

Although climate strongly influenced spatial variation in Zika epidemic metrics, climate was not a strong predictor of Zika epidemic dynamics through time [22,29,37]. We hypothesize that week to week variation in transmission within provinces may be too stochastic to predict based on climate alone. For many of the provinces included in this study, climate rarely fell outside the suitable range for disease transmission, potentially indicating that given suitable climate conditions, non-climate factors such as reporting protocols, underlying immunity, transportation, and infrastructure may generate stochasticity that drives local temporal trends in Zika transmission. Another possible limitation in our ability to predict week to week variation in transmission could be due to errors in the epidemiological data, as suspected case numbers may differ considerably from laboratory-confirmed cases, and case reporting may change over time (e.g., as Zika awareness increases, people may become more likely to go to the doctor for a fever and doctors may be more likely to diagnose symptoms as Zika rather than another disease with similar symptoms such as dengue).

A surprising result of this study was the mismatch between the theoretically and empirically derived optimal temperature for Zika transmission and the temperatures we found to correspond with the greatest force of infection values during the 2015 to 2017 epidemic [25,62]. In previous studies, mechanistic R_0_ models parameterized with data from laboratory experiments indicated that 29°C is optimal for Zika transmission [27,32]. However, our study demonstrated that the highest average force of infection values are found in provinces with mean temperatures between 20 and 26°C (Table 1), and laboratory studies confirm that Zika virus can be detected in the salivary glands of Aedes aegypti and transmitted at 25°C [63]. There are several possible hypotheses that could explain this discrepancy, including that: 1) provinces with mean temperatures around 29°C already implement vector control; 2) people may have acquired cross-immunity from exposure to other arboviruses [25,64,65]; and/or 3) given suitable temperature for transmission, other factors that can covary with temperature such as land use, urbanization, and socioeconomics may be more important drivers of large epidemics [20].These hypotheses raise important questions about how to reconcile and ground truth theoretical and empirical studies with field data.

The 2015 to 2017 Zika outbreak provided an opportunity to investigate how population size and climate affect epidemic dynamics over time and space in a previously unexposed population. The results provide valuable insight into where Zika is likely to become endemic and how future outbreaks transmitted by *Aedes aegypti* could spread. Specifically, given that provinces with greater populations in their largest cities had longer and more intense outbreaks, regions with larger urban population sizes that also have suitable climate conditions for most of the year are most likely to sustain endemic Zika transmission. However, it is important to carefully consider how these results could apply to other regions, especially in places where Zika may be transmitted by other mosquito species, such as *Aedes africanus*, which exhibit different responses to climate [66]. Zika epidemics in Latin America were characteristically different across countries: some regions experienced large epidemics that spread quickly while others experienced low levels of sustained transmission (Figure 2). We would expect similar epidemic metrics to characterize other mosquito transmitted diseases that could emerge and re-emerge in the future, such as o’nyong nyong, Mayaro, and yellow fever [1,19,39–41]. Given this climate-driven geographic variation in epidemics, we would expect climate change to alter patterns in transmission for emerging and re-remerging mosquito-borne diseases in the future.

All data and code are accessible in a github repository: https://github.com/mrc-ide/ZikaModel.

## Author contributions

EAM and JMC conceived of project; MH acquired data and conducted analyses; MH, JMC, and EAM interpreted data and wrote the manuscript.

