## Supplementary Materials for "Climate drives spatial variation in Zika epidemics in Latin America"

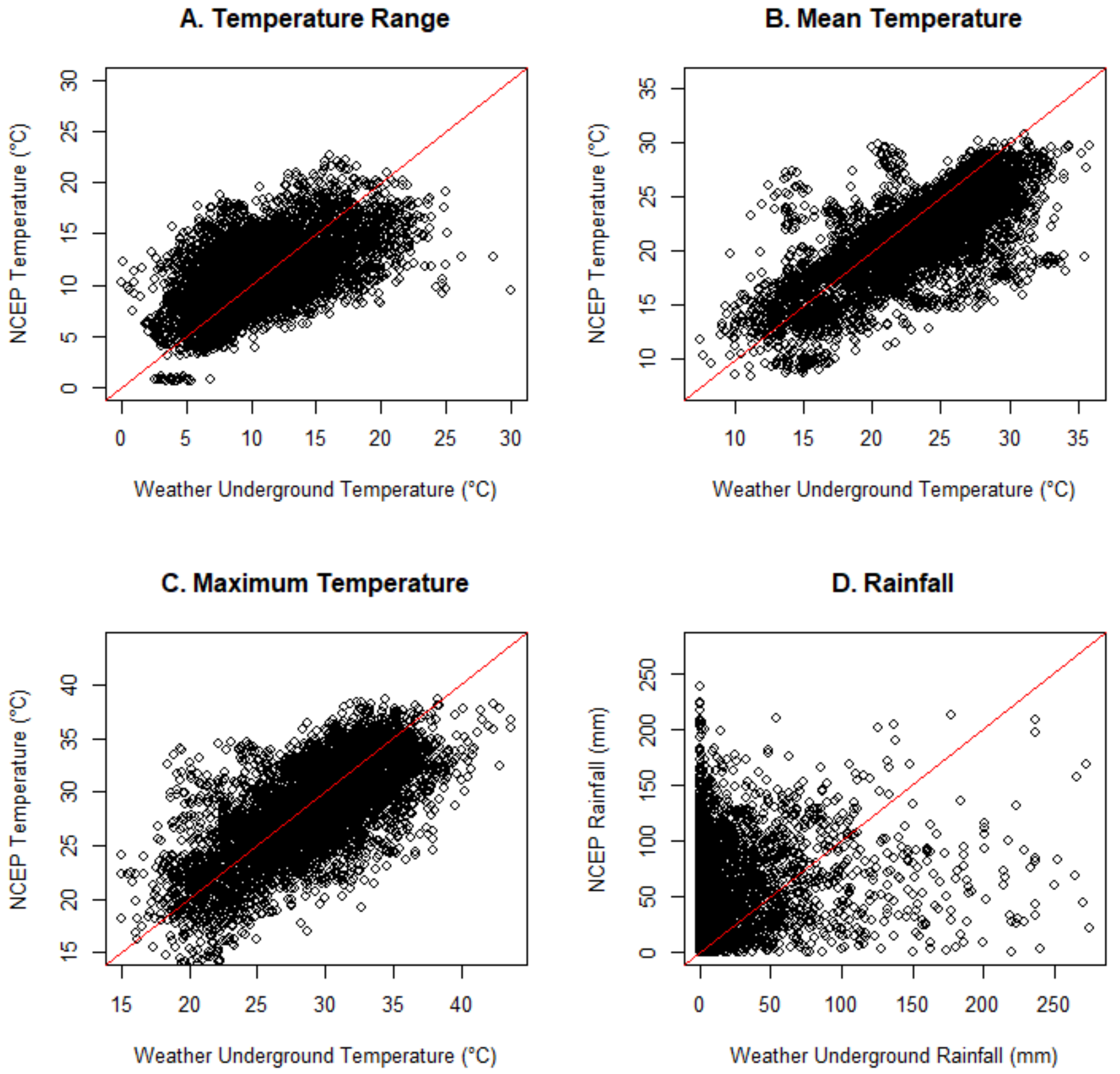


**Figure S1: Scatterplots comparing lagged and aggregated climate variables from the National Centers for Environmental Prediction Reanalysis data (NCEP) and Weather Underground.** NCEP is a gridded global model based on satellite data, available at a 2.5° spatial resolution and six hour temporal resolution [1]. The plots above compare the two sources' values for: A) Temperature range, calculated as the daily difference between the maximum and minimum temperatures observed, aggregated over a three week period and lagged by six weeks from case reporting; B) Average temperature, calculated as the average daily temperature, aggregated over a three week period and lagged by six weeks from case reporting; C) Maximum temperature, calculated as the daily maximum temperature observed, aggregated over a three week period and lagged by six weeks from case reporting; and D) Rainfall, calculated as the daily total precipitation observed, aggregated over a six week period and lagged by three weeks from case reporting. The red line gives y=x, or perfect correspondence between these data. We did not include a plot for humidity, as NCEP measures specific humidity whereas weather underground measures relative humidity, precluding a direct comparison. The NCEP data lacked some important variation observed in the Weather Underground data, as extreme values were less likely to be observed in the NCEP data, likely because of spatial averaging. Rainfall measurements showed a weak relationship with observed rainfall from the weather underground station.

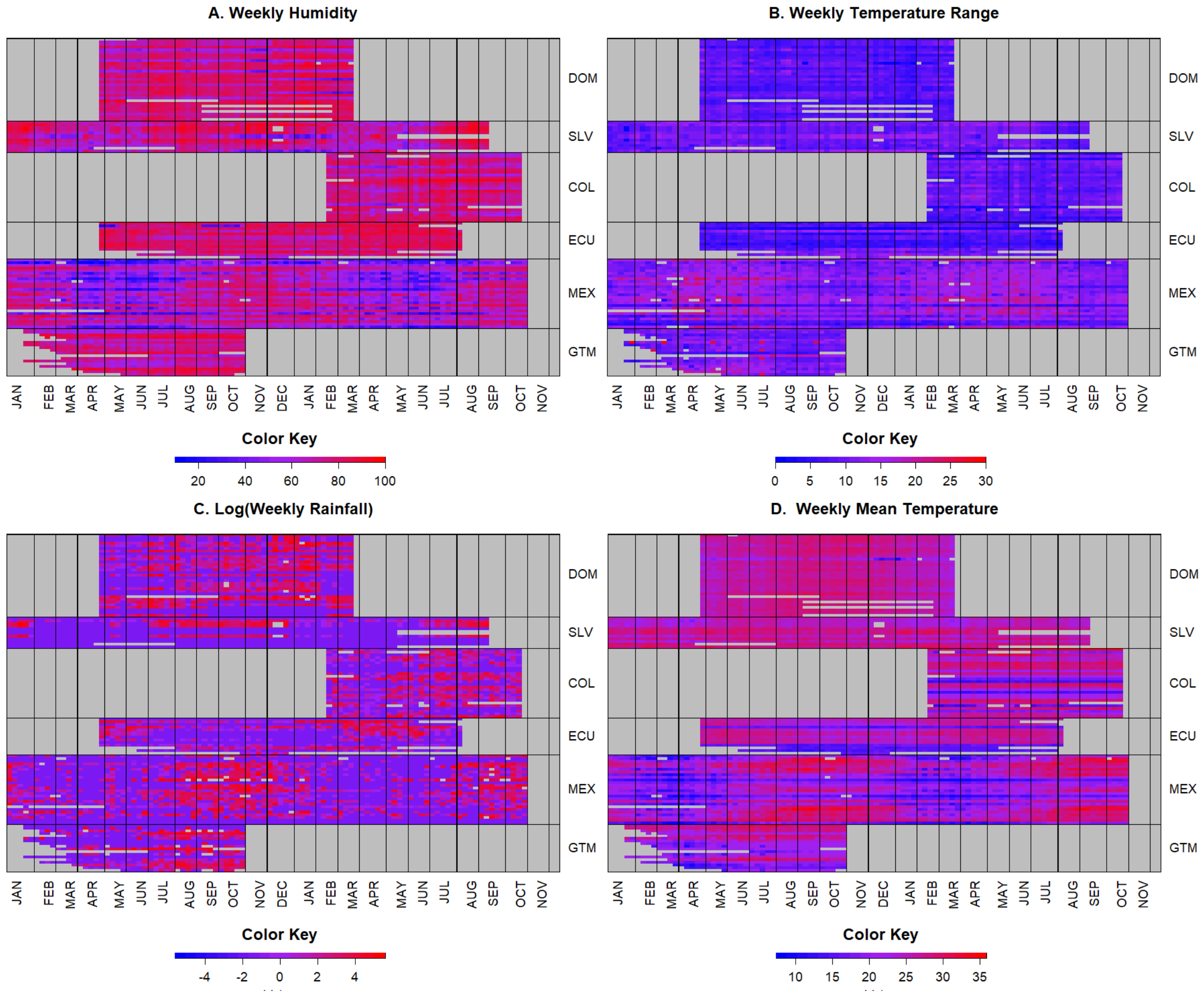


**Figure S2: Heatmaps of Climate Variables Over Time.** Each row corresponds to a single province over time, with lines separating the following countries: DOM = Dominican Republic, SLV = El Salvador, COL = Colombia, ECU = Ecuador, MEX = Mexico, GTM = Guatemala. Gray shading indicates no available data. Red indicates high values and blue indicates low values for appropriately lagged and aggregated climate variables, with panels for (A) humidity, (B) temperature range, (C) log of total rainfall, and (D) mean temperature.

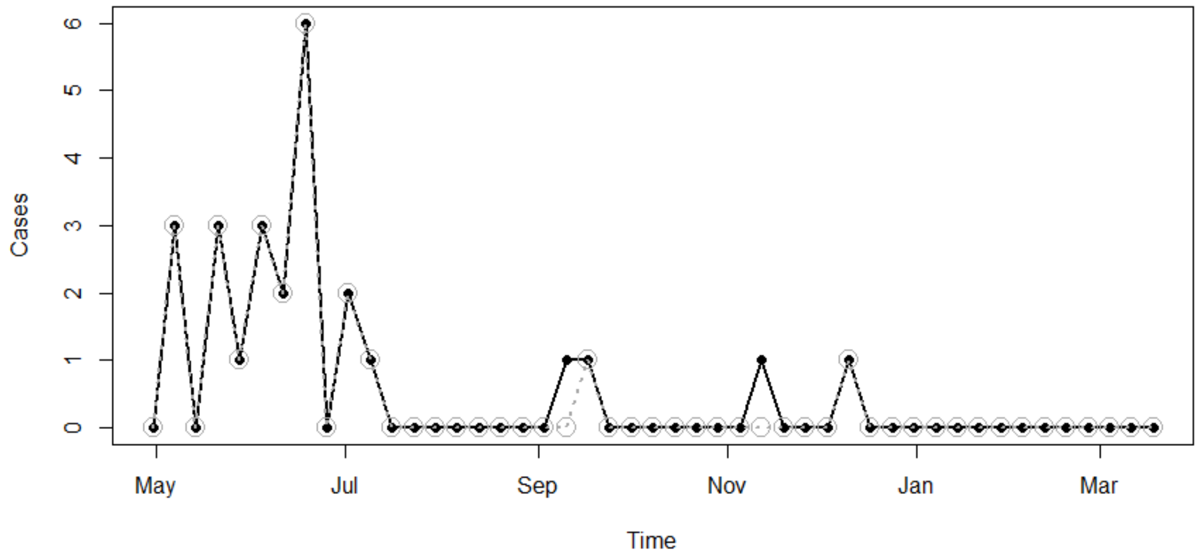


**Figure S3: Sample time series of tSIR Model Fit.** Observed cases (closed points) and predicted cases based on the tSIR model (open circles) plotted over time between May 2016 and March 2017 for a randomly selected province (San Pedro de Macorís, Dominican Republic). Actual cases are given by the solid black line, while predicted cases are given by the dashed gray line. Imported cases, defined as new cases reported in a week with no effective infectious individuals (i.e., no cases reported in the preceding five weeks) are indicated by weeks with zero predicted cases and nonzero actual cases. In weeks with no effective infectious individuals, the transmission parameter $\beta_t$ is set to zero, meaning that the predicted new cases are also zero. In weeks with local transmission (i.e., weeks with effective infectious individuals) there is perfect correspondence between observed cases and predicted cases from the tSIR model because predicted cases were calculated using a linear regression between observed cases (omitting imported cases) and predicted cases. There were 217 imported cases out of a total of 7,109 observations.

‘

**Table S1: Backward elimination for the spatiotemporal hurdle model (step 1): Probability of weekly local transmission, P($\beta_t$>0).** The first row corresponds to the full model; the best fit model, based on the highest adjusted $R^2$ value, is shown in bold. The values in the columns corresponding to climate covariates indicate coefficient values with standard errors reported in parentheses; the statistical significance of coefficients are indicated by asterisks (*=p≤0.05; **=p≤0.01; ***=p≤0.001). Cells containing dashes indicate coefficients that were not included in a particular model. Temperature, humidity, and range correspond to mean temperature, mean humidity, and mean temperature range (difference between the daily maximum and minimum temperatures observed) over a three-week period, lagged by six weeks from the week of case reporting. Temperature$^2$ corresponds to squared temperature over a three-week period, lagged by six weeks from the week of case reporting. Rainfall corresponds to cumulative rainfall over a six-week period, lagged by three weeks from the week of case reporting.

| Model | Temperature | Temperature$^2$ | Humidity | Rainfall | Range | Adj-$R^2$ |
|---|---|---|---|---|---|---|
| P($\beta_t$>0)~ humidity+ temperature+ temperature$^2$+ rainfall+ range | -0.12 (0.15) | 0.18 (0.14) | 0.06 (0.03)* | 0.01 (0.01) | -0.0 (0.03) | 0.029 |
| P($\beta_t$>0)~ humidity+ temperature+ temperature$^2$+ rainfall | -0.12 (0.15) | 0.18 (0.14) | 0.06 (0.02)** | 0.01 (0.01) | - | 0.029 |
| P($\beta_t$>0)~ humidity+ temperature+ temperature$^2$ | -0.11 (0.14) | 0.17 (0.14) | 0.06 (0.02)** | - | - | 0.030 |
| **P($\beta_t$>0)~ humidity+ temperature** | **0.07 (0.03)*** | **-** | **0.06 (0.02)**** | **-** | **-** | **0.037** |
| P($\beta_t$>0)~ humidity | - | - | 0.05 (0.02)* | - | - | 0.023 |

**Table S2: Backward elimination for the spatiotemporal hurdle model (step 2): Intensity of logged weekly force of infection, ln($\beta_t$), given the presence of local transmission.** The first row corresponds to the full model; the best fit model, based on the highest adjusted $R^2$ value, is shown in bold. The values in the columns corresponding to climate covariates indicate coefficient values with standard errors reported in parentheses; the statistical significance of coefficients are indicated by asterisks (*=p≤0.05; **=p≤0.01; ***=p≤0.001). Cells containing dashes indicate coefficients that were not included in a particular model. Temperature, humidity, and range correspond to mean temperature, mean humidity, and mean temperature range (difference between the daily maximum and minimum temperatures observed) over a three-week period, lagged by six weeks from the week of case reporting. Temperature$^2$ corresponds to squared temperature over a three-week period, lagged by six weeks from the week of case reporting. Rainfall corresponds to cumulative rainfall over a six-week period, lagged by three weeks from the week of case reporting.

| Model | Temperature | Temperature$^2$ | Humidity | Rainfall | Range | ΔAdj-$R^2$ |
|---|---|---|---|---|---|---|
| ln(βt)~ humidity+ temperature+ rainfall+ range+ temperature$^2$ | -0.22 (0.26) | 0.16 (0.26) | -0.05 (0.04) | 0.03 (0.03) | 0.04 (0.04) | $9.60 \times 10^{-3}$ |
| ln(βt)~ humidity+ temperature+ rainfall+ range | -0.06 (0.04) | - | -0.05 (0.04) | 0.03 (0.03) | 0.04 (0.04) | $9.13 \times 10^{-3}$ |
| **ln(βt)~ humidity+ temperature+ rainfall** | **-0.07 (0.03)*** | **-** | **-0.08 (0.03)*** | **0.04 (0.03)** | **-** | **$9.64 \times 10^{-3}$** |
| ln(βt)~ humidity+ temperature | -0.07 (0.03)* | - | -0.07 (0.04)* | - | - | $7.99 \times 10^{-3}$ |
| ln(βt)~ humidity | -0.05(0.03) | - | - | - | - | $3.75 \times 10^{-3}$ |

**Table S3: Alternative spatiotemporal panel model with two way fixed effect.** We used the Hausman test to determine whether a random effects or fixed effects model was more appropriate for the spatiotemporal panel models [2]. For both step of the hurdle model, we failed to reject the null hypothesis that the individual and time effects are uncorrelated with the climate covariates concluding that random effects models are more appropriate (weekly local transmission: p=0.947; intensity of weekly force of infection: p=0.370). Hausman testing was conducted in R statistical software version 3.4.3 using the phtest function in the plm package [3]. Compared to the two way random effect panel model described in the manuscript, the two way fixed effect model includes a positive nonlinear effect of temperature, and a positive linear effect of humidity (rather than negative linear). The second step of this model includes a negative nonlinear effect of temperature, as well as a positive linear effect of range, without effects of humidity or rainfall. The values in the columns corresponding to climate covariates indicate coefficient values with standard errors reported in parentheses; the statistical significance of coefficients are indicated by asterisks (*=p≤0.05; **=p≤0.01; ***=p≤0.001).

| Model | Temperature | Temperature$^2$ | Humidity | Rainfall | Range |
|---|---|---|---|---|---|
| $P(\beta_t>0)$~ temperature+ temperature$^2$+ humidity+ rainfall | -0.11 (0.05)* | 0.18 (0.05)*** | 0.06 (0.01)*** | 0.01 (0.01)* | - |
| $\ln(\beta t)$~ temperature+ temperature$^2$+ range | 0.24 (0.34) | -0.36 (0.33) | - | - | 0.06 (0.04) |

**Table S4: Alternative spatiotemporal model with fixed effect of month as a seasonal control.**

The McFadden Pseudo $R^2$ for the first step of this model is 0.078 and the Nagelkerke Pseudo $R^2$ for the second step of this model is 0.044, indicating a poor fit, supporting our hypothesis in the main text that week to week variation in transmission is too stochastic to predict based on climate alone. For both steps of the model, the effect size of temperature is greater than that of the panel linear model. The model of weekly local transmission includes a negative nonlinear effect of temperature, whereas the panel model discussed in the main manuscript includes a linear effect of temperature. Compared to the panel model of weekly local transmission, this model includes an additional effect of range. Compared to the panel spatiotemporal model of intensity of logged weekly force of infection, this model includes an additional effect of rain and a positive nonlinear effect of temperature. The values in the columns corresponding to climate covariates indicate coefficient values with standard errors reported in parentheses; the statistical significance of coefficients are indicated by asterisks (*=p≤0.05; **=p≤0.01; ***=p≤0.001).

| Model | Temperature | Temperature$^2$ | Humidity | Rainfall | Range |
|---|---|---|---|---|---|
| $P(\beta_t>0)$~ temperature+ temperature$^2$+ humidity+ range | 2.06 (0.30)*** | -1.60 (0.29)*** | 0.53 (0.04)*** | - | 0.19 (0.04)*** |
| $\ln(\beta t)$~ temperature+ temperature$^2$+ humidity+ rainfall | -0.44 (0.19)* | 0.35 (0.19) | -0.08 (0.02)*** | 0.05 (0.02)* | - |

**Table S5: Alternative spatiotemporal model with nested fixed effect of month in country as a seasonal control.** The McFadden Pseudo $R^2$ for the first step of this model is 0.229 and the Nagelkerke Pseudo $R^2$ for the second step of this model is 0.181, indicating a poor fit and supporting our hypothesis in the main text that week to week variation in transmission is too stochastic to predict based on climate alone. The effect size of temperature is greater than that of the panel linear model for the presence of weekly local transmission model. The model of weekly local transmission includes a nonlinear effect of temperature, whereas the panel model discussed in the main manuscript includes a linear effect of temperature. Compared to the panel model of weekly local transmission, this model includes an additional effect of range and rainfall. Compared to the panel spatiotemporal model of intensity of weekly force of infection, the nested fixed effect of month in country model does not include an effect of rainfall. The values in the columns corresponding to climate covariates indicate coefficient values with standard errors reported in parentheses; the statistical significance of coefficients are indicated by asterisks (*=p≤0.05; **=p≤0.01; ***=p≤0.001).

| Model | Temp | $Temp^2$ | Hum | Rain | Range |
|---|---|---|---|---|---|
| $P(\beta_t>0)$~ temp+$temp^2$ +hum+rain +range | 1.63 (0.33)*** | -1.38 (0.32)*** | 0.30 (0.05)*** | 0.10 (0.03)*** | 0.09 (0.05) |
| ln(βt)~ temp +hum | -0.05 (0.02)* | - | -0.04 (0.02) | - | - |

**Table S6: Backward elimination for the spatial model of the probability of local transmission in a given province (any week where $\beta_t$>0).** The first row corresponds to the best fit model, based on the Akaike information criterion (AIC). The models are sorted based on increasing AIC value, with the ΔAIC column giving the difference in AIC between the given model and the best fit model. The values in the columns corresponding to climate covariates indicate coefficient values with standard errors reported in parentheses; the statistical significance of coefficients are indicated by asterisks (*=p≤0.05; **=p≤0.01; ***=p≤0.001). Cells containing dashes indicate coefficients that were not included in a particular model. Climate covariates are averaged over the course of the epidemiological data. Temperature, humidity, and range correspond to mean temperature, mean humidity, and mean temperature range (difference between the daily maximum and minimum temperatures observed) over a three-week period, lagged by six weeks from the week of case reporting. Temperature$^2$ corresponds to squared temperature over a three-week period, lagged by six weeks from the week of case reporting. Rainfall corresponds to cumulative rainfall over a six-week period, lagged by three weeks from the week of case reporting. City pop corresponds to the population of the most populated city in each province. The Nakagawa marginal pseudo $R^2$ for the best fit model is 0.281 and the conditional pseudo $R^2$ is 0.599.

| Model | Temperature | Temperature$^2$ | Humidity | Rainfall | Range | City Pop | ΔAIC |
|---|---|---|---|---|---|---|---|
| P(local trans)~ humidity+ temperature+ city pop+ range | 0.49 (0.30) | - | 1.15 (0.48)* | -0.07 (0.45) | - | 0.82 (0.55) | 0 |
| P(local trans)~ humidity+ temperature+ city pop | 0.44 (0.29) | - | 0.94 (0.44)* | 0.08 (0.42) | - | - | 0.492 |
| P(local trans)~ humidity+ temperature+ city pop+ range+rainfall | 0.88 (0.40)* | - | 1.66 (0.62)** | -0.227 (0.45) | 0.99 (0.62) | 1.14 (0.73) | 1.673 |
| P(local trans)~ humidity+ temperature | 0.44 (0.29) | - | 0.96 (0.42)* | - | - | - | 2.821 |
| P(local trans)~ humidity | - | - | 0.92 (0.40)* | - | - | - | 3.319 |
| P(local trans)~ humidity+ temperature+ city pop+ range+rainfall +temperature$^2$ | 2.18 (3.64) | -1.32 (3.69) | 1.66 (0.62)** | -0.30 (0.46) | 1.01 (0.62) | 1.15 (0.73) | 3.528 |

**Table S7: Backward elimination for the spatial model of logged maximum force of infection for each province with local transmission.** The first row corresponds to the best fit model, based on the Akaike information criterion (AIC). The models are sorted based on increasing AIC value, with the ΔAIC column giving the difference in AIC between the given model and the best fit model. The values in the columns corresponding to climate covariates indicate coefficient values with standard errors reported in parentheses; the statistical significance of coefficients are indicated by asterisks (*=p≤0.05; **=p≤0.01; ***=p≤0.001). Cells containing dashes indicate coefficients that were not included in a particular model. Climate covariates are averaged over the course of the epidemiological data. Temperature, humidity, and range correspond to mean temperature, mean humidity, and mean temperature range (difference between the daily maximum and minimum temperatures observed) over a three-week period, lagged by six weeks from the week of case reporting. Temperature$^2$ corresponds to squared temperature over a three-week period, lagged by six weeks from the week of case reporting. Rainfall corresponds to cumulative rainfall over a six-week period, lagged by three weeks from the week of case reporting. City pop corresponds to the population of the most populated city in each province. The Nagelkerke pseudo $R^2$ for the best fit model is 0.534.

| Model | Temperature | Temperature$^2$ | Humidity | Rainfall | Range | City Pop | ΔAIC |
|---|---|---|---|---|---|---|---|
| ln(max(βt))~ temperature+ temperature$^2$ | 2.44 (0.86)** | -2.30 (0.85)** | - | - | - | - | 0 |
| ln(max(βt))~ temperature+ temperature$^2$+ rainfall | 2.46 (0.86)** | -2.34 (0.86)** | - | -0.08 (0.09) | - | - | 1.142 |
| ln(max(βt))~ temperature+ temperature$^2$+ rainfall+range | 2.37 (0.87)** | -2.28 (0.86)** | - | -0.08 (0.09) | -0.09 (0.12) | - | 2.533 |
| ln(max(βt))~ temperature+ temperature$^2$+ rainfall+range+ humidity | 2.37 (0.87)** | -2.29 (0.97)** | -0.02 (0.11) | -0.08 (0.09) | -0.10 (0.14) | - | 4.488 |
| ln(max(βt))~ temperature | 0.15 (0.09) | - | - | - | - | - | 5.520 |
| ln(max(βt))~ temperature+ temperature$^2$+ rainfall+range+ humidity+ city pop | 2.37 (0.88)** | -2.29 (0.87)* | -0.02 (0.11) | -0.08 (0.09) | -0.10 (0.14) | - 0.01(0.10) | 6.478 |

**Table S8: Backward elimination for the spatial model of logged mean force of infection for each province with local transmission.** The first row corresponds to the best fit model, based on the Akaike information criterion (AIC). The models are sorted based on increasing AIC value, with the ΔAIC column giving the difference in AIC between the given model and the best fit model. The values in the columns corresponding to climate covariates indicate coefficient values with standard errors reported in parentheses; the statistical significance of coefficients are indicated by asterisks (*=p≤0.05; **=p≤0.01; ***=p≤0.001). Cells containing dashes indicate coefficients that were not included in a particular model. Climate covariates are averaged over the course of the epidemiological data. Temperature, humidity, and range correspond to mean temperature, mean humidity, and mean temperature range (difference between the daily maximum and minimum temperatures observed) over a three-week period, lagged by six weeks from the week of case reporting. Temperature$^2$ corresponds to squared temperature over a three-week period, lagged by six weeks from the week of case reporting. Rainfall corresponds to cumulative rainfall over a six-week period, lagged by three weeks from the week of case reporting. City pop corresponds to the population of the most populated city in each province. The Nagelkerke pseudo $R^2$ for the best fit model is 0.596.

| Model | Temp | Temp$^2$ | Hum | Rain | Range | City.pop | ΔAIC |
|---|---|---|---|---|---|---|---|
| ln(mean(βt))~ temperature+ temperature$^2$+ city pop | 2.75 (0.82)*** | -2.78 (0.81)*** | - | - | - | 0.19 (0.09)* | 0 |
| ln(mean(βt))~ temperature+ temperature$^2$+ city pop+ range | 2.78 (0.82)*** | -2.70 (0.81)** | - | -0.02 (0.08) | -0.15 (0.11) | 0.21 (0.09)* | 0.185 |
| ln(mean(βt))~ temperature+ temperature$^2$+ city pop+ range+rainfall | 2.78 (0.82)*** | -2.70 (0.81)** | | -0.02 (0.08) | -0.15 (0.11) | 0.21 (0.09)* | 2.148 |
| ln(mean(βt))~ temperature+ temperature$^2$ | 2.75 (0.82)** | -2.65 (0.82)** | - | - | - | - | 2.774 |
| ln(mean(βt))~ temperature+ temperature$^2$+ city pop+ range+rainfall+ humidity | 2.78 (0.82)** | -2.69 (0.81)** | 0.02 (0.10) | -0.02 (0.08) | -0.14 (0.13) | 0.21 (0.09) | 4.124 |
| ln(mean(βt))~ temperature | 0.11 (0.09) | - | - | - | - | - | 11.568 |

**Table S9: Backward elimination for the spatial model of cumulative cases reported.** The first row corresponds to the best fit model, based on the Akaike information criterion (AIC). The models are sorted based on increasing AIC value, with the ΔAIC column giving the difference in AIC between the given model and the best fit model. The values in the columns corresponding to climate covariates indicate coefficient values with standard errors reported in parentheses; the statistical significance of coefficients are indicated by asterisks (*=p≤0.05; **=p≤0.01; ***=p≤0.001). Cells containing dashes indicate coefficients that were not included in a particular model. Climate covariates are averaged over the course of the epidemiological data. Temperature, humidity, and range correspond to mean temperature, mean humidity, and mean temperature range (difference between the daily maximum and minimum temperatures observed) over a three-week period, lagged by six weeks from the week of case reporting. Temperature$^2$ corresponds to squared temperature over a three-week period, lagged by six weeks from the week of case reporting. Rainfall corresponds to cumulative rainfall over a six-week period, lagged by three weeks from the week of case reporting. City pop corresponds to the population of the most populated city in each province. The Nagelkerke pseudo $R^2$ for the best fit model is 0.330.

| Model | Temperature | Temperature$^2$ | Humidity | Rainfall | Range | City Pop | ΔAIC |
|---|---|---|---|---|---|---|---|
| cumulative cases~ city pop+ temperature+ temperature$^2$+ humidity | 1.69 (0.86) | -1.68 (0.86) | 0.16 (0.09) | - | - | 0.21 (0.09)* | 0 |
| cumulative cases~ city pop+ temperature+ temperature$^2$+ humidity+ rainfall | 1.66 (0.86) | -1.63 (0.86) | 0.12 (0.10) | 0.09 (0.09) | - | 0.20 (0.09)* | 0.714 |
| cumulative cases~ city pop+ temperature+ temperature$^2$ | 1.85 (0.86)* | -1.84 (0.86)* | - | - | - | 0.20 (0.09)* | 1.068 |
| cumulative cases~ city pop | - | - | - | - | - | 0.20 (0.09)* | 1.893 |
| cumulative cases~ city pop+ temperature+ temperature$^2$+ humidity+ rainfall+range | 1.57 (0.88) | -1.57 (0.87) | 0.09 (0.11) | 0.09 (0.09) | -0.08 (0.13) | 0.20 (0.09)* | 2.272 |
| cumulative cases~ city pop+ temperature | 0.02 (0.09) | - | - | - | - | 0.20 (0.09)* | 3.848 |

**Table S10: Backward elimination for the spatial model of mean weekly cases reported in weeks that cases were reported.** The first row corresponds to the best fit model, based on the Akaike information criterion (AIC). The models are sorted based on increasing AIC value, with the ΔAIC column giving the difference in AIC between the given model and the best fit model. The values in the columns corresponding to climate covariates indicate coefficient values with standard errors reported in parentheses; the statistical significance of coefficients are indicated by asterisks (*=p≤0.05; **=p≤0.01; ***=p≤0.001). Cells containing dashes indicate coefficients that were not included in a particular model. Climate covariates are averaged over the course of the epidemiological data. Temperature, humidity, and range correspond to mean temperature, mean humidity, and mean temperature range (difference between the daily maximum and minimum temperatures observed) over a three-week period, lagged by six weeks from the week of case reporting. Temperature$^2$ corresponds to squared temperature over a three-week period, lagged by six weeks from the week of case reporting. Rainfall corresponds to cumulative rainfall over a six-week period, lagged by three weeks from the week of case reporting. City pop corresponds to the population of the most populated city in each province. The Nagelkerke pseudo $R^2$ for the best fit model is 0.348.

| Model | Temperature | Temperature$^2$ | Humidity | Rainfall | Range | City Pop | ΔAIC |
|---|---|---|---|---|---|---|---|
| mean weekly cases~ city pop+ temperature+ temperature$^2$+ rainfall | 1.69 (0.85)* | -1.59 (0.85) | - | 0.14 (0.08) | - | 0.35 (0.09)*** | 0 |
| mean weekly cases~ city pop+ temperature+ temperature$^2$+ rainfall+humidity | 1.59 (0.85) | -1.50 (0.85) | 0.12 (0.10) | 0.11 (0.08) | - | 0.36 (0.09)*** | 0.212 |
| mean weekly cases~ city pop+ temperature+ temperature$^2$ | 1.79 (0.85)* | -1.72 (0.85)* | - | - | - | 0.36 (0.09)*** | 1.416 |
| mean weekly cases~ city pop+ temperature+ temperature$^2$+ rainfall+humidity+ range | 1.49 (0.86) | -1.43 (0.86) | 0.09 (0.11) | 0.11 (0.08) | -0.09(0.13) | 0.37 (0.09)*** | 1.647 |
| mean weekly cases~ city pop | - | - | - | - | - | 0.35 (0.09)*** | 2.708 |
| mean weekly cases~ city pop+ temperature | 0.08 (0.09) | - | - | - | - | 0.36 (0.09)*** | 3.704 |
